# A machine learning approach to predicting short-term mortality risk in patients starting chemotherapy

**DOI:** 10.1101/204081

**Authors:** Aymen A. Elfiky, Maximilian J. Pany, Ravi B. Parikh, Ziad Obermeyer

## Abstract

**Background:** Cancer patients who die soon after starting chemotherapy incur costs of treatment without benefits. Accurately predicting mortality risk from chemotherapy is important, but few patient data-driven tools exist. We sought to create and validate a machine learning model predicting mortality for patients starting new chemotherapy.

**Methods:** We obtained electronic health records for patients treated at a large cancer center (26,946 patients; 51,774 new regimens) over 2004-14, linked to Social Security data for date of death. The model was derived using 2004-11 data, and performance measured on non-overlapping 2012-14 data.

**Findings:** 30-day mortality from chemotherapy start was 2.1%. Common cancers included breast (21.1%), colorectal (19.3%), and lung (18.0%). Model predictions were accurate for all patients (AUC 0.94). Predictions for patients starting palliative chemotherapy (46.6% of regimens), for whom prognosis is particularly important, remained highly accurate (AUC 0.92). To illustrate model discrimination, we ranked patients initiating palliative chemotherapy by model-predicted mortality risk, and calculated observed mortality by risk decile. 30-day mortality in the highest-risk decile was 22.6%; in the lowest-risk decile, no patients died. Predictions remained accurate across all primary cancers, stages, and chemotherapies—even for clinical trial regimens that first appeared in years after the model was trained (AUC 0.94). The model also performed well for prediction of 180-day mortality (AUC 0.87; mortality 74.8% in the highest risk decile vs. 0.2% in the lowest). Predictions were more accurate than data from randomized trials of individual chemotherapies, or SEER estimates.

**Interpretation:** A machine learning algorithm accurately predicted short-term mortality in patients starting chemotherapy using EHR data. Further research is necessary to determine generalizability and the feasibility of applying this algorithm in clinical settings.

## INTRODUCTION

Chemotherapy lowers the risk of cancer recurrence in early-stage cancers, and can improve survival and symptoms in later stage disease. Balancing these benefits against chemotherapy’s considerable risks is challenging. There is growing evidence that chemotherapy is started too often, too late in the cancer disease trajectory,^1–4^ and many patients die soon after initiating treatment. These patients experience burdensome symptoms and financial costs, without many of the potential benefits of chemotherapy.^5^ National organizations now track the fraction of patients who die within two weeks of receiving chemotherapy as a marker of poor quality of care,^6,7^ and this number has been rising rapidly.^1,8^

A key factor underlying these trends is the difficulty of accurately predicting the risk of serious adverse events, especially death, when initiating chemotherapy. Side effects of chemotherapy are variable, and the influence of comorbidities is complex, all making the risk calculus challenging.^9–12,13^ Cognitive biases also lead to underestimation of risk of death,^14–15^ particularly in patients with metastatic cancer^16,17^ who often believe that their disease is curable.^18,19^ Physicians themselves are notoriously bad at estimating prognosis in patients with cancer,^20,21^ and overly optimistic estimates can influence patients’ chemotherapy decisions.^26–31^

Currently, doctors may use randomized trial data to estimate mortality for individual regimens, or online tools based on Surveillance, Epidemiology, and End Results (SEER) data to obtain mortality risk by age, sex, and primary cancer.^14,28^ While informative, these tools provide mortality estimates for broad populations of patients, and their relevance to individual decisions is unclear. Individualized decision support tools do exist,^29^ but require a substantial investment of time and resources to collect and enter data not readily available in existing records. There is considerable enthusiasm for the role of advanced algorithms to improve prediction, by drawing on the rich data stored in electronic health records (EHRs).^30^ However, there is little evidence that such algorithms can provide meaningful inputs to clinical decision making, in cancer or elsewhere.

Here we develop and validate a machine learning algorithm to predict near-term mortality risk in patients starting new chemotherapy regimens. New chemotherapy is a critical event in the cancer disease trajectory, and can serve as a ‘pause point’ to weigh difficult questions. Objective predictions of short-term mortality at this time could be useful to doctors and patients in several ways. First and foremost, estimated likelihood of serious adverse events is an important input to discussions of risks and benefits of treatments, particularly for patients undergoing palliative chemotherapy.^31–34^ Accurate forecasts could also help guide important decisions for patients around family and financial arrangements. Finally, patients at high mortality risk could be prompted to complete advance care planning processes or offered palliative care consultation.

## METHODS

### Study Population

We obtained EHR data for all cancer patients receiving chemotherapy at the Dana-Farber/Brigham and Women’s Cancer Center (DF/BWCC) from 2004–2014. We determined date of death by linking to the Social Security Administration’s Death Master File. We classified patients by primary cancer and presence of distant-stage disease, determined using registry data (for patients diagnosed at DF/BWCC) and International Classification of Diseases (ICD) codes for metastases (for patients not diagnosed at DF/BWCC or who did not have registry data; and to identify progression to distant-stage disease in those previously diagnosed at DF/BWCC).^35^ While diagnosis codes have limitations for determination of cancer stage, they are generally believed to reliably identify presence or absence of distant-stage disease.^36^ The institutional review boards of Dana-Farber Cancer Institute and Partners HealthCare approved this study and granted a waiver of informed consent from study participants.

### Statistical Analysis

#### Dataset

Our primary outcome was death within 30 days of starting new systemic chemotherapy regimens. Secondary outcomes were 30-day mortality in pre-specified subgroups of interest (described below) as well as overall 180-day mortality. We constructed our dataset at the patient–chemotherapy regimen level, such that each observation was a new regimen.

#### Model performance

Machine learning models have the potential to ‘overfit’, or produce overly optimistic estimates of model performance based on spurious correlations in development data. We thus report results only in an independent validation set, which played no role in model development; as such, overfitting would only lead to poorer model performance in the validation set. Specifically, we used data from 2004–2011 for model derivation, and data from 2012–2014 for model validation. Importantly, while our dataset was at the patient–chemotherapy regimen level, we randomly assigned patients— not observations—to the derivation or validation sets, since observations describing different chemotherapy regimens in the same patient were not independent. As such, no patient appeared in both sets.

We report area under the receiver operating characteristic curve (AUC)^37^ with 95% confidence intervals,^38^ overall and in subgroups of interest, notably age, sex, race, distant-stage disease, individual primary cancers, chemotherapy lines and regimens, and chemotherapy intent (palliative vs. curative, identified by the treating physician, via an EHR flag). To benchmark against existing prognostic models, we obtained one-year mortality estimates from large randomized trials of specific chemotherapies, and from the Surveillance, Epidemiology, and End Results (SEER) program.

#### Predictors

To transform raw EHR data into variables usable in a prediction model, we first pulled all data from the one-year period ending the day before chemotherapy initiation (we did not drop patients based on absence of data over this period). Raw data were aggregated into 23,641 potential predictors, in the following categories: demographics, prescribed medications, comorbidities and other grouped^35^ ICD-9 diagnoses, procedures,^35^ care utilization, vital signs, laboratory results, and terms derived from physician notes. For each potential predictor, we created two variables, the sum of related EHR entries over two time periods: 0-1 months (recent) and 1-12 months (baseline) prior to chemotherapy initiation. This strategy is outlined in more detail elsewhere.^39^ We also included a variable indexing how many lines of chemotherapy the patient had in total prior to the current regimen. No data on the current regimen itself (agent, intent, etc.) were used in the predictive model. We dropped variables missing in over 99% of the development sample, leaving 5,390 predictors in the model.

#### Algorithm

We used high-dimensional statistical techniques designed to handle large sets of correlated predictors, specifically gradient boosted trees: a linear combination of decision trees similar to those used to derive many clinical decision rules^40^ (R package: xgboost).^41^ We used 4-fold cross-validation in the development sample to choose model parameters (*e.g.*, number of trees, variables per tree). The model was configured to produce individual-level probabilities of 30-day mortality. More details are available in the Supplemental Methods.

#### Missing values

Each split of each tree in the model (*e.g.*, a split on sex) had a ‘default’: the value (*e.g.*, male or female) that occurred more frequently in the training data. Observations with missing values for a given variable were assigned to the default side of the split. This was effectively a split-specific, probabilistic imputation function that allowed us to avoid dropping observations missing data.

#### Model parameters

Given the complexity of the model, a succinct summary of its parameters was challenging. We attempted to do so by breaking down model predictions into the linear contributions of individual variables, using a decomposition of model predictions on all predictors (in the development sample). We calculated the (linear) sum of squares for each individual variable included in the machine learning model, and interpreted the residual sum of squares as the contribution of non-linear terms and interactions used by the model. Since our model used over 5000 predictors, we chose to report only a small selection, specifically those that most explained model variance, and those identified as predictors of mortality in prior studies.^29,42,43^

## RESULTS

### Study Population

We identified 26,946 patients initiating 51,774 discrete chemotherapy regimens over 2004–2014; 59.4% had distant-stage disease. **Table 1** shows baseline patient characteristics at time of chemotherapy initiation. The most common chemotherapy regimens (derivation and validation sets) were carboplatin–paclitaxel (n=4042), gemcitabine (n=2185), and albumin-bound paclitaxel (n=1985); 3.4% of the validation set (n=523) received chemotherapy regimens that first appeared in 2012 or later, and thus did not appear in the derivation set, including experimental, non-FDA-approved agents (2.3%; n=343).

**Table 1.** Baseline patient characteristics of model derivation and validation sets.

| *Variable* | **Sample** |  | *Difference* |
| --- | --- | --- | --- |
|  | *Derivation* | *Validation* |  |
| Patients, n | 17,832 | 9,114 | 8,718 |
| Chemo regimens, n | 36,007 | 15,767 | 20,240 |
| Age, mean (95% CI) | 58.7 (58.5 to 58.9) | 60.7 (60.5 to 61) | -2.1 (-2.4 to -1.7) |
| Female, % (95% CI) | 61.1 (60.4 to 61.9) | 60.7 (59.7 to 61.7) | 0.5 (-0.7 to 1.7) |
| White, % (95% CI) | 86.9 (86.4 to 87.4) | 88.3 (87.7 to 89) | -1.5 (-2.3 to -0.7) |
| IP visit, % (95% CI) | 43.9 (43.2 to 44.6) | 38 (37 to 38.9) | 6 (4.7 to 7.2) |
| Gagne score, median (IQR) | 2 (1 to 6) | 1 (1 to 6) | -1 (0 to 0) <sup>a</sup> |
| Cancer type, % (95% CI) |  |  |  |
| Breast | 23.6 (23 to 24.3) | 21.1 (20.2 to 21.9) | 2.5 (1.5 to 3.6) |
| Colon and Rectum | 17.6 (17.1 to 18.2) | 19.3 (18.5 to 20.2) | -1.7 (-2.7 to -0.7) |
| Lung and Bronchus | 17.7 (17.2 to 18.3) | 18 (17.2 to 18.8) | -0.3 (-1.3 to 0.7) |
| Hematological | 7.1 (6.7 to 7.5) | 8.4 (7.8 to 8.9) | -1.3 (-2 to -0.6) |
| Ovary | 6.7 (6.3 to 7) | 7.7 (7.1 to 8.2) | -1 (-1.7 to -0.3) |
| Other | 27.8 (27.1 to 28.4) | 25.8 (24.9 to 26.7) | 2 (0.9 to 3.1) |
| Chemo beyond first line, % (95% CI) | 49 (48.3 to 49.7) | 42.4 (41.4 to 43.4) | 6.6 (5.3 to 7.8) |
| Intent of chemotherapy % (95% CI) <sup>b</sup> |  |  |  |
| Curative | 15.7 (15.3 to 16.1) | 26.8 (26.1 to 27.5) | -11.1 (-11.9 to -10.3) |
| Palliative | 33.8 (33.3 to 34.3) | 46.6 (45.8 to 47.3) | -12.7 (-13.7 to -11.8) |

**Note:**
<sup>a</sup> Median (IQR) is reported with difference calculated by quantile regression. 95% CI shows difference between the IQRs of the derivation and validation cohorts.
<sup>b</sup> Intent of chemotherapy does not add up to 100% due to missing values, resulting in unknown intent.

There were several significant differences between the 2004–11 derivation set and the 2012–14 validation set, including age at initiation, race, primary cancer, and prior chemotherapy beyond the first line. Such differences between derivation and validation sets were expected, and indeed intentional: a validation set drawn from later years of data was chosen to reflect the constant evolution of cancer epidemiology and treatment. This made the prediction task more difficult, since algorithms trained on past data cannot always perform well in the future,^44^ but accurately represented the difficulties algorithms face in evolving real-world settings.

### Model performance

Among patients in the validation set, overall 30-day mortality was 2.1%, and 3.1% among those initiating chemotherapy with palliative intent. The model performed accurately for predicting 30-day mortality for all patients, irrespective of chemotherapy intent (AUC: 0.94; 95% CI, 0.93 to 0.95). It also performed well when restricting to patients receiving palliative chemotherapy (AUC: 0.92; 95% CI, 0.91 to 0.94). To illustrate the concrete implications of this, we used model predictions to individually rank patients by 30-day mortality risk, a commonly used way of stratifying risk groups.^37^ 30-day mortality in the highest decile of predicted risk was 22.6%, while in the lowest risk decile, not a single patient died.

**Figure 1** shows observed survival over the 180 days after palliative chemotherapy initiation by decile of model-predicted 30-day mortality risk. 180-day mortality was higher overall, at 18.4%, but model-predicted 30-day mortality was also an accurate predictor of 180-day mortality (AUC 0.87). 180-day mortality in the highest risk decile was 74.8%, vs. 0.2% in the lowest risk decile.

**Figure 1.**
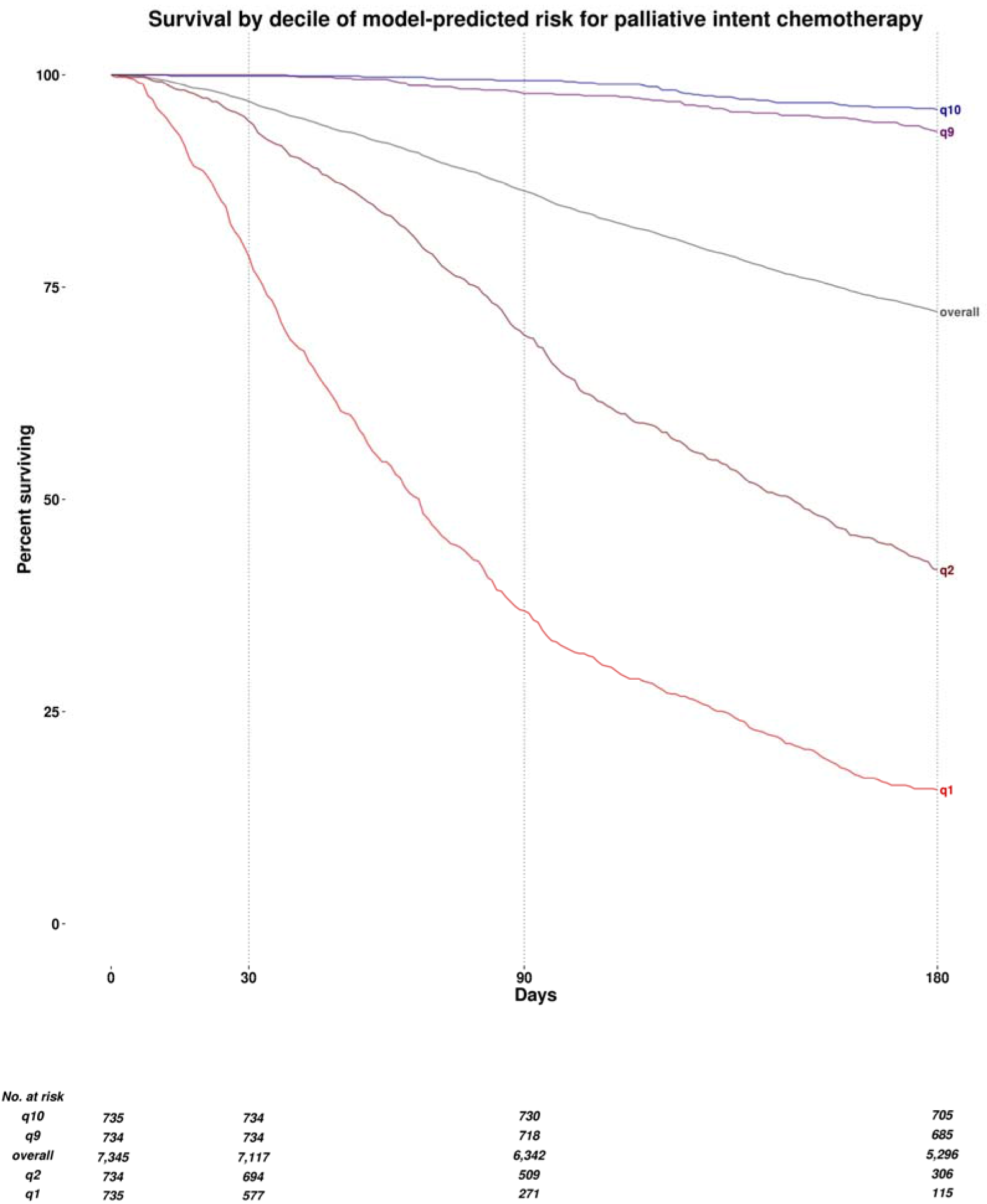
Observed survival over the 180 days from the initiation of palliative chemotherapy, by decile of model-predicted mortality risk. *q1* denotes the highest predicted risk decile, *q2* the second highest, *q9* the second lowest, and *q10* the lowest. *Overall* denotes overall mean survival in all patients, irrespective of model-predicted risk.

**Table 2** shows model performance for predicting 30-day mortality in additional patient subgroups of interest. The model performed equally well across many kinds of primary cancers, demographic groups, and chemotherapy regimens. In distant-stage disease (average 30-day mortality: 2.9%), 30-day mortality in the highest risk decile was 22.7%, vs. 0.0% in the lowest decile (AUC: 0.94; 95% CI, 0.93 to 0.95). Strikingly, predictions were accurate even for experimental new clinical trial regimens first observed over 2012–14 (AUC: 0.94; 95% CI, 0.88 to 1.0)—i.e., regimens that first appeared in years of data to which the model was not exposed in the training process.

**Table 2.** Model performance in selected subgroups.

| Subgroup | AUC (95% CI) |  |
| --- | --- | --- |
|  | 30-day mortality | 180-day mortality |
| *Intent of chemotherapy* |  |  |
| Curative | 0.981 (0.970 to 0.992) | 0.892 (0.872 to 0.912) |
| Palliative | 0.924 (0.910 to 0.939) | 0.827 (0.817 to 0.838) |
| *Stage* |  |  |
| Distant stage | 0.938 (0.926 to 0.951) | 0.852 (0.842 to 0.861) |
| Non-distant stage | 0.936 (0.916 to 0.955) | 0.874 (0.862 to 0.887) |
| *Cancer* |  |  |
| Breast | 0.970 (0.956 to 0.983) | 0.939 (0.928 to 0.951) |
| Colon and Rectum | 0.924 (0.898 to 0.949) | 0.843 (0.827 to 0.860) |
| Lung and Bronchus | 0.916 (0.884 to 0.949) | 0.820 (0.803 to 0.838) |
| Ovary | 0.956 (0.922 to 0.989) | 0.895 (0.871 to 0.918) |
| Hematological | 0.943 (0.902 to 0.983) | 0.819 (0.782 to 0.856) |
| Head and Neck | 0.970 (0.933 to 1.000) | 0.839 (0.798 to 0.879) |
| Cervix Uteri | 0.955 (0.906 to 1.000) | 0.888 (0.847 to 0.929) |
| Prostate | 0.936 (0.849 to 1.000) | 0.805 (0.751 to 0.858) |
| *Chemotherapy regimen* |  |  |
| Carbo/paclitaxel | 0.940 (0.859 to 1.000) | 0.846 (0.813 to 0.880) |
| FOLFOX | 0.952 (0.923 to 0.982) | 0.863 (0.833 to 0.893) |
| Gemcitabine | 0.865 (0.789 to 0.942) | 0.992 (0.982 to 1.000) |
| Pemetrexed/carboplatin | 0.960 (0.939 to 0.981) | 0.807 (0.769 to 0.845) |
| Paclitaxel | 0.989 (0.972 to 1.000) | 0.738 (0.686 to 0.791) |
| Novel clinical trial agents <sup>a</sup> | 0.942 (0.882 to 1.000) | 0.870 (0.817 to 0.923) |
| *Line of chemotherapy* |  |  |
| First line | 0.941 (0.925 to 0.956) | 0.865 (0.854 to 0.875) |
| Subsequent line | 0.938 (0.924 to 0.952) | 0.864 (0.854 to 0.874) |
| *Year <sup>b</sup>* |  |  |
| 2012 | 0.951 (0.935 to 0.966) | 0.887 (0.875 to 0.899) |
| 2013 | 0.935 (0.914 to 0.955) | 0.869 (0.857 to 0.882) |
| 2014 | 0.928 (0.908 to 0.947) | 0.837 (0.824 to 0.851) |
| *Age group* |  |  |
| 21-40 | 0.963 (0.939 to 0.988) | 0.901 (0.874 to 0.928) |
| 41-60 | 0.950 (0.935 to 0.966) | 0.893 (0.883 to 0.903) |
| 61-80 | 0.936 (0.920 to 0.951) | 0.850 (0.839 to 0.861) |
| 81-100 | 0.876 (0.806 to 0.947) | 0.803 (0.768 to 0.838) |
| *Gender* |  |  |
| Female | 0.948 (0.935 to 0.962) | 0.886 (0.877 to 0.894) |
| Male | 0.927 (0.910 to 0.944) | 0.840 (0.828 to 0.852) |
| *Race* |  |  |
| White | 0.940 (0.929 to 0.951) | 0.870 (0.862 to 0.877) |
| Non-white | 0.941 (0.909 to 0.972) | 0.870 (0.850 to 0.891) |
| *Insurance* |  |  |
| Private | 0.949 (0.937 to 0.962) | 0.886 (0.878 to 0.895) |
| Medicare | 0.938 (0.911 to 0.965) | 0.854 (0.837 to 0.872) |
| Medicaid | 0.945 (0.914 to 0.976) | 0.824 (0.784 to 0.865) |
| Self-pay | 0.932 (0.851 to 1.000) | 0.821 (0.753 to 0.888) |

<sup>a</sup> Subgroup of patients receiving new clinical trial regimens first observed over 2012–14, i.e., years of data to which the model was not exposed in the training process.
<sup>b</sup> Observations in the validation set occurring in 2011 (n = 717) were reassigned to 2012 for the purpose of this table.

A key question is whether model predictions are accurate enough to be useful across a range of primary cancers, stages of disease, lines of chemotherapy—scenarios whose prognoses vary widely. Table 2 thus also presents measures of overall predictive accuracy for first line (AUC 0.94 for 30-day mortality, 0.87 for 180-day mortality) vs. later lines (AUC 0.94 for 30-day mortality, 0.86 for 180-day mortality) of chemotherapy. eTable 1 presents extended results on accuracy for 30- and 180-day mortality across lung, colorectal, breast, and prostate cancers by stage and line of chemotherapy.

### Comparisons to other prognostic estimates

We compared model performance to two external sources of mortality predictions, focusing on patients with distant-stage disease for whom prognostic estimates are most valuable.

First, we obtained mortality data from four randomized trials of treatments for colorectal adenocarcinoma, non-small cell lung adenocarcinoma, small cell lung carcinoma, and squamous cell carcinoma of the head and neck.^45–48^ **Figure 2a** shows observed one-year mortality (the only mortality outcome reported consistently) for patients on these regimens in our validation sample, compared to: (1) point estimates of average one-year mortality from relevant clinical trials, and (2) quintiles of estimated mortality risk (30-day) from our model. The overall AUC for RCT mean estimates was 0.555 (95% CI, 0.513 to 0.598), compared to 0.771 (95% CI, 0.735 to 0.808) for our individual-level model-based estimates for these same patients.

**Figure 2.**
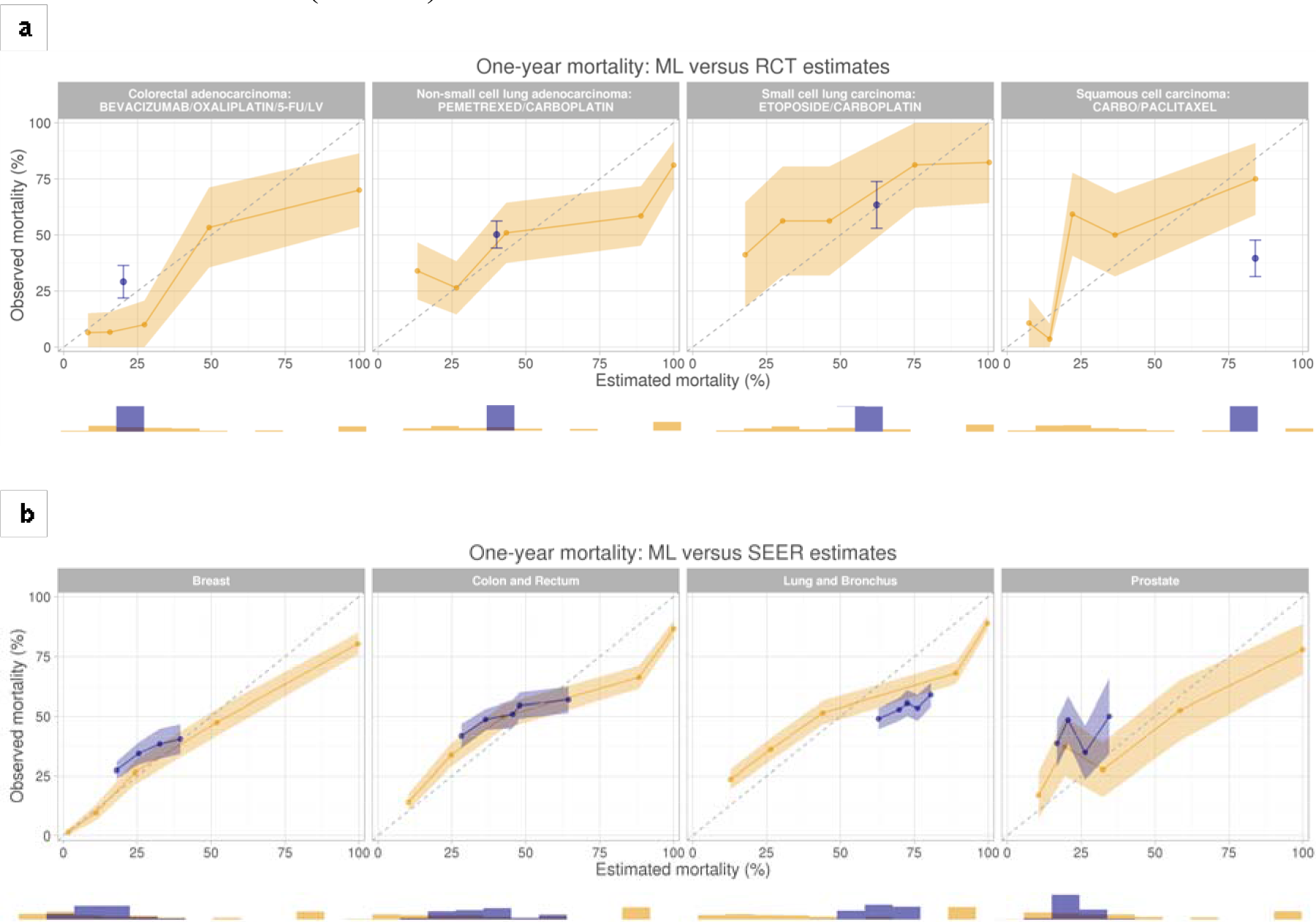
Observed one-year mortality (y-axis) from start of chemotherapy for patients with specific combinations of distant-stage cancers and chemotherapies, shown against mortality predictions from three sources (x-axis). The 45° dotted line denotes equivalence of observed and estimated mortality. Orange lines, with shaded 95% confidence intervals, show mean observed one-year mortality against quintiles of estimated one-year mortality based on our model-predicted 30-day mortality risk. Blue lines, with shaded 95% confidence intervals, show observed one-year mortality against predictions from two other sources. In **Panel (a)**, we show mean one-year mortality and 95% CI (blue dots and bars) from individual randomized trials. Of note, trial data provide only single point estimates for mortality, based on average mortality in the trial, leading to one point estimate on the graphs. In **Panel (b)**, we show mean observed one-year mortality against quantiles of estimated one-year mortality risk from the National Cancer Institute’s Surveillance, Epidemiology, and End Results (SEER) program, estimated using cancer type, age, sex, and race. Histograms show the distribution of individual estimates of mortality risk, colored by source of estimate: orange for our model predictions, blue for RCT estimates (Panel a) and SEER estimates (Panel b).

We also compared our mortality predictions to age-, sex-, race-, and cancer-specific mortality estimates from SEER, restricting to patients with advanced-stage cancers of lung and bronchus, colon and rectum, breast, and prostate to maximize comparability in populations. **Figure 2b** shows that our estimates were far more accurate (AUC: 0.810; 95% CI, 0.799 to 0.822) than SEER estimates (AUC: 0.600; 95% CI, 0.585 to 0.615) for predicting one-year mortality. Further details on construction of RCT and SEER estimates are in the Supplemental Methods and eTable 2, and more detailed comparisons for subgroups are available in eTable 3.

**Table 3.** Selected variables by risk decile and model variance explained.

|  |  | Variable mean, by risk decile |  |  | Model<br>variance<br>explained |
| --- | --- | --- | --- | --- | --- |
| Variable |  | Top | Median | Bottom |  |
| Cancer | Brain and other NS | 6.7 | 1.6 | 0.0 | 1.94% |
| Demographics | Age | 62.3 | 62.1 | 51.9 | 1.30% |
|  | Female | 56.4 | 60.9 | 86.9 | 1.17% |
|  | Black | 3.8 | 3.6 | 3.4 | 0.07% |
| Comorbidity score |  | 5.14 | 3.44 | 2.01 | 0.03% |
| Prior diagnoses | Ascites | 0.31 | 0.07 | 0.01 | 0.39% |
|  | Mouth disorder | 0.02 | 0.01 | 0.01 | 0.25% |
|  | Nausea and vomiting | 0.23 | 0.09 | 0.01 | 0.18% |
|  | Lower respiratory disorders | 2.10 | 1.36 | 0.16 | 0.03% |
|  | Secondary malignancy | 5.57 | 1.69 | 0.36 | 0.01% |
|  | Failure to thrive | 0.05 | 0.01 | 0.00 | 0.01% |
|  | Nutritional disorders | 2.53 | 1.63 | 0.54 | 0.00% |
|  | Malaise and fatigue | 0.22 | 0.08 | 0.02 | 0.00% |
| Medications | Corticosteroids | 0.53 | 0.00 | 0.00 | 0.15% |
|  | Opioids | 0.29 | 0.00 | 0.00 | 0.05% |
|  | Anxiolytics | 0.60 | 0.24 | 0.18 | 0.02% |
|  | Cathartic | 0.54 | 0.00 | 0.00 | 0.00% |
| Vital signs <sup>a</sup> | Pulse, max (year) | 106.1 | 95.7 | 87.1 | 0.37% |
|  | Weight, max (year) | 79.5 | 80.2 | 76.8 | 0.11% |
|  | Weight, standard deviation (year) | 3.1 | 2.1 | 1.3 | 0.06% |
|  | Pulse, min (month) | 83.4 | 72.8 | 63.0 | 0.01% |
|  | Weight change <sup>b</sup> | -3.1 | -1.0 | 0.1 | 0.00% |
| Labs <sup>a</sup> | C-reactive protein, max | 93.9 | 65.6 | 2.2 | 0.19% |
|  | SGPT, max | 75.9 | 57.3 | 24.3 | 0.07% |
|  | SGOT, max | 73.7 | 54.2 | 23.8 | 0.09% |
|  | White blood cell count, max | 13.9 | 12.4 | 9.8 | 0.03% |
|  | Alkaline phosphatase, max | 199.5 | 128.7 | 76.5 | 0.02% |
|  | Lymphocytes, mean | 1.0 | 1.3 | 1.8 | 0.00% |
|  | Platelets, mean | 251.8 | 241.4 | 265.1 | 0.00% |
| Other | Ejection fraction | 54.4 | 48.0 | 51.9 | 0.01% |
| Total, linear terms |  |  |  |  | 13.6 |
| Total, nonlinear terms and interactions |  |  |  |  | 86.4 |

<sup>a</sup> “Year” and “month” denote the value of the variable (minimum, maximum, or standard deviation, as noted) over the 12 months and 1 month prior to chemotherapy initiation, respectively
<sup>b</sup> Denotes the change in weight from the average value over the 12 months prior to chemotherapy initiation to the average value over the 1 month prior to chemotherapy initiation
<sup>c</sup> Denotes the value of the variable (minimum, maximum, as noted) over the 12 months prior to chemotherapy initiation.

### Key predictors

**Table 3** shows the distribution of key predictor variables used in the prediction model across risk deciles, as well as the proportion of model variance explained linearly by each variable. In general, key predictors of mortality identified in the literature were markedly different for patients in the highest vs. lowest model-predicted risk deciles: for example, summed comorbidity score^43^, age^42^, failure to thrive, heart rate, and certain laboratory data (e.g. C-reactive protein, white blood cell count, alkaline phosphatase).^29^ But importantly, while these differences were striking, no one variable explained more than 2% of model predictions on its own. Indeed, the majority of variation in the predictions (86.4%) was not a linear function of any one predictor, indicating that the tree-based model relied heavily on complex non-linear functional forms and interactions among variables.

## DISCUSSION

A machine learning model based on single-center EHR data accurately estimated individual mortality risk at the time of chemotherapy initiation. The model performed well across a range of cancer types, race, sex, and other demographic variables. Mortality estimates were accurate for palliative as well as curative chemotherapy regimens, for early-and distant-stage patients, and even for patients treated with clinical trial regimens introduced in years after the model was trained. Our model dramatically outperformed estimates from randomized trials and SEER data, both of which are routinely used by clinicians for quantitative risk predictions.

It is notable that this model was able to predict mortality with considerable accuracy despite lacking genetic sequencing data, cancer-specific biomarkers, or indeed any detailed information about cancers beyond EHR data. This underscores the fact that common clinical data elements contained within an EHR—e.g. symptoms, comorbidities, prescribed medications, diagnostic tests—contain surprising amounts of signal for predicting key outcomes in cancer patients.

Algorithmic predictions such as ours could be useful at several points along the care continuum. This could include providing accurate predictions of mortality risk to a provider or tumor board at the ‘point of decision,’ or fostering shared decision making between patient and provider at the ‘point of care.’ Short-term mortality risk predictions could help clinicians identify patients unlikely to benefit from chemotherapy beyond 30 days, and those who may benefit from early palliative care referral, advance care planning, and prompting to get financial and family affairs in order. For patients receiving systemic chemotherapy, predictable 30-day mortality may even be a useful quality indicator of avoidable treatment-associated harm.^49^

Importantly, while machine learning algorithms require significant computing infrastructure to construct, once derived, they can be applied using only the computing power available on a personal computer or smartphone. This greatly facilitates potential integration into existing clinical systems. While our algorithm was developed using a single institution’s data, its data inputs are representative of what is generally available in structured format in EHRs, including ICD and procedure codes, medications, etc. Thus, there are no technical barriers to implementing this or similar algorithms in any organization’s clinical data to independently validate predictive power out of sample. To this end, code for our algorithm will be made publicly available (at http://labsysmed.org/wp-content/uploads/2017/02/ChemoMortalityAnalysis.rtf).

This study has several limitations. Our model was built on data from patients treated with chemotherapy, meaning that it answers the question: what is the mortality risk of a a patient starting chemotherapy today? This can be highly useful, since prognostic information informs many important decisions for patients and doctors at a critical point in the disease trajectory. But it also has several important caveats. First, predictions are unlikely to be accurate for untreated patients, meaning the model cannot answer the related question: what is the effect of chemotherapy on mortality risk? Second, our treated sample reflects the particular decisions around chemotherapy made by doctors and patients in our training dataset. Patients who could or would have started chemotherapy, but for some reason did not, would not be included, which can bias the sample. But—for better or worse—the direction of this bias is predictable: prevailing treatment decisions are generally aggressive. In our sample, 62.4% of patients with distant-stage disease received chemotherapy, and evidence suggests that physicians in a wide range of settings overestimate survival, and overuse chemotherapy. Thus, to the extent that there is bias in our dataset, it leads to the inclusion—not exclusion—of marginal patients, who otherwise might not have received chemotherapy. As a result, we believe this bias did not substantially distort validity. If such an algorithm were deployed in a real-world setting, periodic re-training of the model (*e.g.*, each year or quarter) would ensure that model predictions reflected contemporaneous chemotherapy decision-making. This would address changing selection into treatment over time, and update the model to reflect broader changes in patient populations and chemotherapy technology.

While we took pains to quantify predictive accuracy in an independent, recent validation set, the only way to truly validate such a model is prospectively. A model trained on pre-2012 data may lose accuracy as novel tumor diagnostics and therapies come online, although the accuracy of predictions for patients starting novel chemotherapies was encouraging in this regard. In addition, this is a single-institution study. Further validation is required using cohorts from different institutions. EHR data contain a multitude of biases introduced by physician behavior, institutional idiosyncrasies, and software platforms, among other limitations. These can significantly affect its adaptability and relevance to different care settings.

In conclusion, our machine learning model accurately predicted mortality risk in patients at the time of chemotherapy initiation. While we are optimistic that accurate prognostic tools such as this could help to promote value-driven oncology care, the ideal next step would be a randomized trial of algorithmic estimates at the point of care. To be useful, predictive models must improve decision-making in the real world. Thus rigorous evaluation of predictions’ impact on outcomes is the gold standard test—but one that is often neglected in the literature, which focuses primarily on measuring predictive accuracy rather than real outcomes.

